# Negative affective states are not detected in rats following an intravenous self-administration regimen leading to incubation of oxycodone craving

**DOI:** 10.64898/2026.04.06.716594

**Authors:** Amanda M. Wunsch, Kimberley A. Mount, Asia Guzman, Alex B. Kawa, Jonathan G. Westlake, Hayley M. Kuhn, Madelyn M. Beutler, Marina E. Wolf

## Abstract

In rats, cue-induced opioid craving intensifies (incubates) during abstinence from opioid self-administration and then remains high for a prolonged period. The prolonged plateau models persistent vulnerability to cue-induced craving and relapse in humans recovering from opioid use disorder. However, a very significant contributor to relapse vulnerability in these individuals is the presence of negative affective states that can persist for months to years, far beyond physical dependence. The goal of this study was to determine if the incubation of craving model recapitulates this aspect of relapse vulnerability. We began by comparing rats trained to self-administer oxycodone using a regimen leading to persistent elevation of cue-induced craving (6 h/d x 10 d) and rats trained to self-administer saline. We assessed somatic withdrawal signs in early abstinence and conducted behavioral tests modeling negative affect (open field, social preference, sucrose preference, and elevated plus maze) in late abstinence. Some somatic withdrawal signs were greater in oxycodone rats on abstinence day (AD)1, but cumulative scores did not differ between groups on AD1-3. On AD41-46, no group differences were found in behavioral tests modeling negative affect. To compare early and late abstinence periods, a second cohort of rats self-administered saline and oxycodone and then received two cue-induced seeking tests (AD1 and AD40; oxycodone rats exhibited incubation of craving) and two series of negative affect tests (AD2-7 and AD41-48). While some time-dependent changes in affect were observed within each group, they were suggestive of reduced anxiety-like behavior in oxycodone rats. Finally, because rats are single-housed during our incubation studies, we compared drug-naïve rats after 8-9 weeks of single vs pair housing and found no difference in behavioral tests modeling negative affect. We conclude that the persistence of elevated cue-induced craving observed after a standard opioid incubation regimen is not accompanied by negative affective states, probably due to lower drug intake during the intravenous regimen compared to non-contingent escalating dose regimens typically used to study withdrawal signs. This does not negate the utility of the incubation model for studying cue-induced opioid craving and its neurobiological basis.

## 1. Introduction

In people with opioid use disorder (OUD), craving is strongly correlated with likelihood of relapse (1), and this vulnerability to relapse persists during protracted abstinence (2). Cue-induced drug craving and relapse can be studied using the incubation of craving model. In rats, cue-induced seeking for drugs of abuse (stimulants, nicotine, alcohol, and opioids) progressively intensifies (incubates) during forced or voluntary abstinence from drug self-administration and then remains high for a prolonged period (3). In humans, incubation has been demonstrated during abstinence from cocaine, methamphetamine, nicotine and alcohol but remains controversial for opioids (3) (see Discussion). Nevertheless, persistence of craving during abstinence, captured by the plateau phase of the incubation model, applies across all drugs. The model’s focus on cue-induced craving is also translationally relevant. Abstinent opioid users exhibit robust motivational bias towards opioid-associated stimuli (4) and such bias predicts heroin relapse (5) and is correlated with craving (6). Thus, the incubation of craving model has translational relevance to certain aspects of OUD.

However, it is well established that negative affective states of opioid abstinence (including loss of motivation for natural rewards, anxiety, dysphoria, and irritability) contribute very significantly to relapse vulnerability; importantly, they persist for months to years in recovering opioid users, far beyond somatic withdrawal signs (7, 8). In rodents, these states have been extensively studied after escalating dose non-contingent opioid regimens leading to physical dependence (9, 10). While physical dependence wanes in days to a week, negative affective states are more persistent and sometimes intensify from 1 to 4-6 weeks of withdrawal (9, 10). Tests that often revealed progressively intensifying negative affective states include social interaction, elevated plus maze (EPM), open field (center time), and tests of motivation for natural rewards (11–20). Far less is known about the persistence of negative affective states following drug self-administration, as most such studies have focused on the early abstinence period (see Discussion).

The goal of this study was to determine if negative affective states occur alongside incubation of opioid craving. We focused on oxycodone, a widely prescribed opioid that is also widely abused (21) and that elicits incubation of craving in rats after forced or voluntary abstinence (3). Negative affective states have mainly been assessed in early abstinence from oxycodone self-administration (see Discussion), but it is important to know if they persist at protracted withdrawal times when incubation is expressed. If so, it would set the stage for future studies to test the interaction of negative affective states and cue-learning in persistent drug craving. To this end we employed open field, social preference test, sucrose preference test, and elevated plus maze (selected based on studies cited in the previous paragraph) after different periods of abstinence from saline or oxycodone self-administration. We used an extended access oxycodone regimen (0.1 mg/kg/infusion x 6 h/day x 10 days) that produced robust incubation in our previous study (22). This oxycodone dose is commonly used in incubation studies across labs [recent examples: (23–25)].

## 2. Materials and Methods

### 2.1 Animals

We obtained rats on a Long Evans background (26 male and 28 female) from our in-house breeding colony and used both sexes in each experiment. Our recent study (22) and most others (see Discussion) have found similar oxycodone incubation in male and female rats. We housed rats 2-3 per cage until the time of jugular catheter surgery or single housing, at which time we individually housed rats. Rats were 7-15 weeks old at the start of the experiment. Rats had free access to food and water and were maintained on a 12 h light/dark cycle. We excluded 3 rats for health-related issues. All procedures were approved by the OHSU Institutional Animal Care and Use Committee and conform to guidelines outlined in the *Guide for the Use and Care of Animals* and *Policy on Humane Care and Use of Laboratory Animals*.

### 2.2 Jugular Catheter Surgery

Rats destined for oxycodone(or saline) self-administration (SA) received jugular catheter implants for drug delivery. We anesthetized rats using isoflurane (5% induction, 2-3% maintenance) and injected them with meloxicam (5 mg/kg, SC) for analgesia. We shaved surgical sites and prepared them with betadine and ethanol scrubs. Jugular catheters consisted of a mesh-reinforced, 22-gauge cannula with an attached 10 cm silastic tube. We placed the exterior cannula on the back, threaded the tube subcutaneously around the shoulder blade, and inserted the tube into the jugular vein. We applied topical antibiotics to the incisions and administered prophylactic cefazolin (30 mg/kg, IV). Rats received 3 d of post-operative meloxicam and recovered from surgery for 5-8 d before the start of drug SA. We flushed catheters 5-7 d per week with heparinized saline (20 IU/mL, IV) until the cessation of drug SA.

### 2.3 Oxycodone SA

Rats self-administered oxycodone (0.1 mg/kg/infusion; NIDA Drug Supply Program) or 0.9% saline for 6 h/d for 10 consecutive days under a fixed ratio 1 schedule as described previously (22). Nose pokes in the active hole triggered the infusion pump and simultaneously activated a 4 s cue light and time out period. Pokes in the inactive hole had no consequence. Following SA, rats remained in their home cage for the duration of forced abstinence except for periods of behavior testing.

### 2.4 Cue-induced seeking

Rats returned to the SA chamber for 1 h. Nose pokes in the previously active hole resulted in presentation of the same cue light previously paired with drug but without drug infusion. Pokes in the inactive hole continued to result in no consequences. The number of pokes in the previously active hole is our measure of cue-induced craving.

### 2.5 Spontaneous somatic withdrawal (SWD) symptoms

Assessments occurred in an acrylic cylinder (30 cm diameter x 50 cm tall) with bedding on the floor. Habituation to the cylinder occurred during two 10 min sessions prior to SA. On abstinence day (AD) 1-7, we placed rats in the cylinder for 10 min while video-recording behavior. An experimenter blind to experimental condition counted the frequency of digging, jumping, rearing, grooming, and teeth chatters. We report SWD symptoms as a total behavior score, which is the sum of the total occurrences of each behavior, as well as reporting each behavior individually. We modified previous procedures (26, 27) to include only behaviors we could reliably detect from videos. SWD scores on only AD1-3 are presented here because no differences from saline controls were detected on later ADs. We recorded rat weight prior to each session.

### 2.6 Open Field Test (OFT)

This test, which can be used to measure anxiety-like behavior (28), was performed in an arena with textured floors (100 x 100 x 45 cm). We placed rats in the center of the arena and allowed them to freely explore for 5 min. ANYmaze software tracked the center of the body and calculated the total distance travelled and the number of entries and time spent in the periphery (10 cm around edge) and center (inner 80 cm^2^). The percentage of time spent in the center of the arena was used as a measure of anxiety-like behavior.

### 2.7 Social Preference Test (SoPT)

This test, commonly used to study social motivation and reward (29), was carried out in an apparatus consisting of 3 chambers separated by walls with spaces that allow rats free access to all chambers. The outer 2 chambers are 50 x 60 cm and the center chamber is 30 x 60 cm. Two wire cups are placed on opposite sides of the apparatus. During habituation, we placed the test rat in the center of the chamber and allowed it to freely explore the entire apparatus for 5 min. During the test phase, we placed a stranger conspecific (same sex as the test rat) under one of the wire cups (location of stranger conspecific was counterbalanced across test subjects). We then placed the test rat in the center of the chamber, and the rat freely explored the apparatus for 10 min. ANYmaze software tracked the center of the rat and calculated the number of entries into and time spent exploring the chamber with the stranger conspecific or the empty cup. We calculated the social preference score as the percentage of time the test rat spent exploring the stranger conspecific divided by total time spent exploring the stranger conspecific and empty cup.

### 2.8 Sucrose Preference Test (SuPT)

We used the SuPT as a measure of anhedonia or depressive-like behavior. We performed a 3 d SuPT protocol as described previously (30) in the rat’s home cage. To habituate the rats to the two-bottle choice paradigm and overcome neophobia, rats had access to two bottles, one containing 1% sucrose solution and one that was empty, for 2 d. Rats did not consume normal water during this period. On day 3, we removed both bottles from the cage, and rats underwent a 4 h water deprivation period to increase motivation to consume liquid during the test period. During the 1 h test, rats had access to one bottle containing 1% sucrose and a second bottle containing water. We gave rats fresh sucrose bottles and switched the location of the sucrose bottle daily. We measured sucrose and water consumption by weighing the bottles before and after the test and calculated sucrose preference as the amount of sucrose consumed over the total amount of sucrose and water consumed.

### 2.9 Elevated Plus Maze (EPM)

This test, widely used to measure anxiety-like behavior (28), was performed in an apparatus in the shape of a plus sign (+) raised 75 cm off the floor, with 2 arms encased by opaque walls (50 x 10 x 60 cm) and 2 open arms without walls (50 x 10 cm). We placed rats in the center of the apparatus and allowed them to freely explore for 5 min. ANYmaze software tracked the center of the body and calculated the number of entries and time spent in the center, closed, and open arms. The percentage of time spent in the open arm relative to the total time spent in the open and closed arms was our measure of anxiety-like behavior.

### 2.10 Statistical Analyses

We used GraphPad Prism to plot data and perform statical analyses. We tested for normal distribution using the Shapiro-Wilk test. Two Way Mixed ANOVAs were used to assess infusions taken during SA (drug and day as factors), nose pokes during both SA and cue-induced seeking tests (hole and drug as factors), and time-dependent changes in spontaneous SWD signs, body weight, active and inactive nose pokes during seeking tests, and OF, SoPT, SuPT, and EPM data (abstinence day and drug as factors). Significant interactions were followed up with uncorrected Fisher’s LSD post hoc tests and significant main effects were followed up using Sidak’s multiple comparison’s test. One Way ANOVA followed by Tukey’s multiple comparisons test or non-parametric Kruskal-Wallis test followed by Dunn’s multiple comparisons test were used to assess OF, SoPT, SuPT, and EPM in single-housed naïve rats and rats that underwent oxycodone or saline self-administration. Unpaired t-tests or nonparametric Mann-Whitney U tests were used to assess average infusions taken during SA or OF, SuPT, SoPT, and EPM data in pair-housed and single-housed rats. Rats of both sexes were used in all experiments, and we observed no obvious differences between males and females for any measure. Because these studies were not sufficiently powered to detect sex differences, data from both sexes are combined. All data are presented as mean ± SEM.

## 3. Results

### 3.1 Spontaneous somatic withdrawal (SWD) symptoms and behaviors indicative of negative affect during abstinence from oxycodone or saline self-administration

The goal of the first experiment was to determine if our oxycodone SA regimen elicits SWD symptoms in early abstinence and/or negative affective states after protracted abstinence (timeline in Fig. 1A). Key ANOVA results are reported here. For full ANOVA results see Table 1 (for Fig. 1 data) or Table S1 (for Fig. S1 data); for post hoc test results see legends to Fig. 1 and Fig. S1.

**Figure 1.**
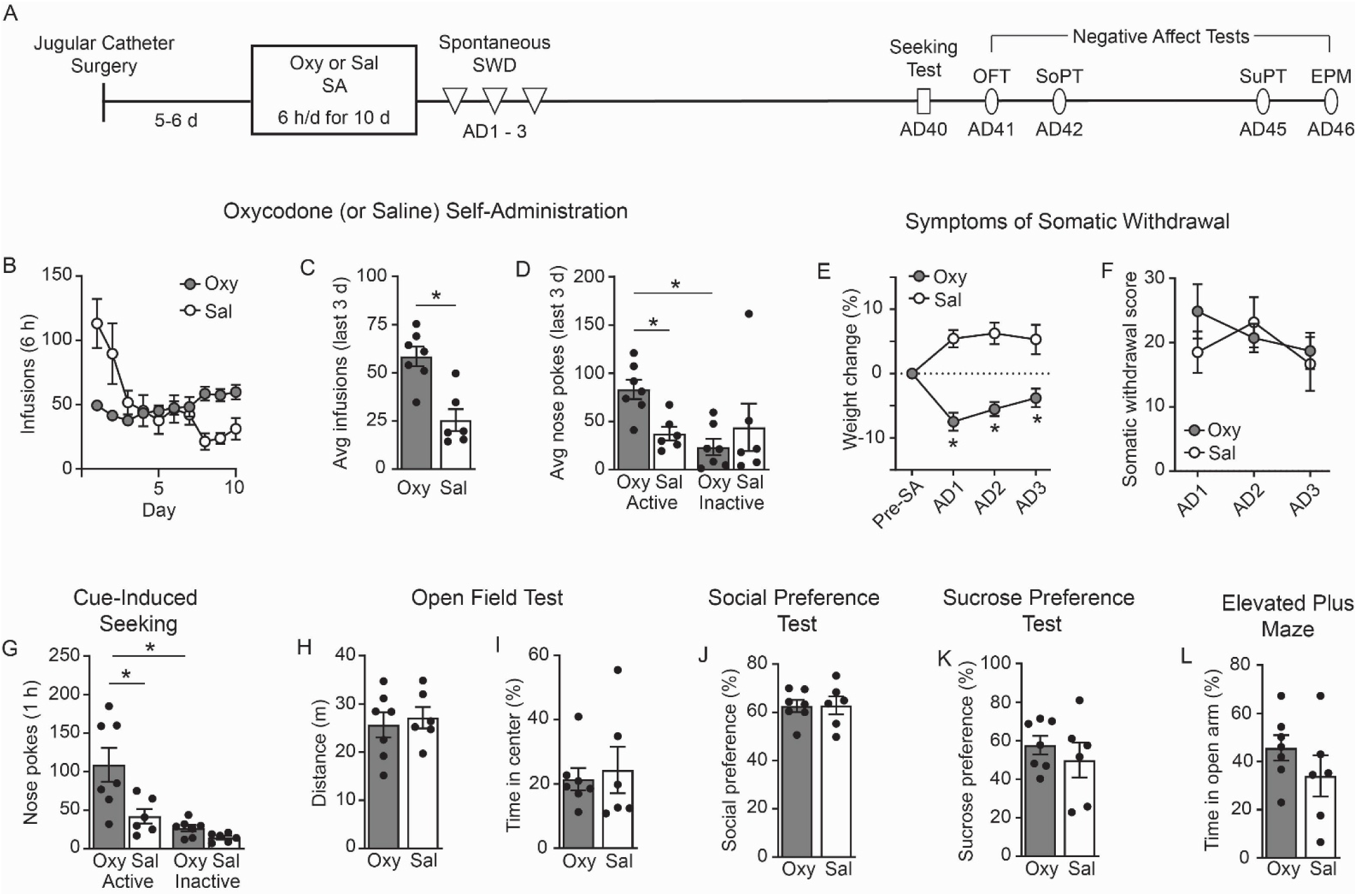
Oxycodone self-administration (SA), symptoms of somatic withdrawal following cessation of oxycodone self-administration, and cue-induced oxycodone craving and negative affective states after 40 days of forced abstinence. A) Rats received jugular catheters and recoveredfor 5-6 days. Rats self-administered oxycodone (Oxy; 0.1 mg/kg/infusion; n=7) or saline (Sal; n=6) for 6 h/d for 10 d. We assessed symptoms of spontaneous somatic withdrawal (SWD) on abstinence day (AD) 1-3 following SA. Rats then underwent a cue-induced seeking test (1 h) on AD40, open field test (OFT) on AD41, social preferencetest (SoPT) on AD42, sucrose habituation on AD43-44, sucrose preference test (SuPT) on AD45, and elevated plus maze (EPM) on AD46. B) The number of infusions of Oxy or Sal that rats self-administered in 6 h across the 10 d of SA. During early days of training, several saline rats took a higher than typical number of infusions but there was no reason for excluding these animals. C) During the last 3 d of SA, Oxy rats took a significantly higher number of infusions than Sal rats (unpaired t-test, p=0.0012). D) During the last 3 d of SA, Oxy rats poked more in the active hole than the inactive hole (post hoc uncorrected Fisher’s LSD, p=0.0094), an effect not observed in Sal rats. Oxy rats poked more in the active hole than Sal rats (post hoc uncorrected Fisher’s LSD, p=0.0271). E) Oxy and Sal rats exhibited different patterns in weight changes on AD1-3 (post hoc Sidak’s multiple comparisons, all p<0.01), with the weight of Oxy rats significantly decreased on AD1-3 compared to their pre-SA weight (post hoc Sidak’s multiple comparisons, all p<0.02). F) The somatic withdrawal score (sum of occurrences of grooming bouts, wet dog shakes, teeth chatters, jumps, and digging bouts in 10 min) did not differ between Oxy and Sal rats on AD1-3. G) Oxy rats poked in the previously active hole significantly more than Sal rats (post hoc Sidak’s multiple comparisons, p=0.0029) and in the inactive hole (post hoc Sidak’s multiple comparisons, p=0.0030) during a cue-induced seeking test. H-I) Oxy and Sal rats travelled the same distance (H) and spent the same percentage of time in the center of the arena (I) during the OFT. J) Oxy and Sal rats spent a similar percentage of time in the chamber with a novel conspecific rat during SoPT. K) Oxy and Sal rats consumed the same percentage of sucrose during the SuPT. L) Oxy and Sal rats spent the same percentage of time in the open arm during EPM. All data presented as mean ± SEM; individual data are shown as closed circles in bar graphs. *p<0.05. Abbreviations: abstinence day (AD), average (avg), elevatedplus maze (EPM), open field test (OFT), oxycodone (Oxy), saline (Sal), self-administration (SA), social preference test (SoPT), somatic withdrawal (SWD), sucrose preference test (SuPT). Full statistical results for Fig. 1 are provided in Table 1. See Fig. S1, Table S1, and Table S2 for related analyses.

**Table 1.** Full statistical results for all data presented in Figures 1, 2 and 3.

| Figure | Statistical Test | F, H, t, or U | p-value |
| --- | --- | --- | --- |
| **Figure 1: Early abstinence somatic withdrawal and late abstinence negative affect** |  |  |  |
| 1B: SA infusions | Two Way Mixed ANOVA |  |  |
| | Drug x Day interaction | $F_{(9,99)}=14.20$ | <0.0001* |
| | Within subject: Day | $F_{(1.572,17.30)}=9.071$ | 0.0034* |
| | Between subject: Drug | $F_{(1,11)}=0.02383$ | 0.8801 |
| 1C: SA avg infusions | Unpaired t-test (Oxy, Sal) | $t_{(11)}=4.348$ | 0.0012* |
| 1D: SA avg pokes | Two Way Mixed ANOVA |  |  |
| | Hole x Drug interaction | $F_{(1,11)}=5.589$ | 0.0375* |
| | Within subject: Hole | $F_{(1,11)}=3.611$ | 0.0839# |
| | Between subject: Drug | $F_{(1,11)}=0.9043$ | 0.3621 |
| 1E: AD1-3 weight | Two Way Mixed ANOVA |  |  |
| | AD x Drug interaction | $F_{(3,33)}=23.90$ | <0.0001* |
| | Within subject: AD | $F_{(1.288,14.17)}=1.625$ | 0.2284 |
| | Between subject: Drug | $F_{(1,11)}=27.83$ | 0.0003* |
| 1F: AD1-3 SWD | Two Way Mixed ANOVA |  |  |
| | AD x Drug interaction | $F_{(2,22)}=1.597$ | 0.2252 |
| | Within subject: AD | $F_{(1.610,17.71)}=1.867$ | 0.1876 |
| | Between subject: Drug | $F_{(1,11)}=0.2444$ | 0.6308 |
| 1G: AD40 seeking | Two Way Mixed ANOVA |  |  |
| | Hole x Drug interaction | $F_{(1,11)}=3.607$ | 0.0841# |
| | Within subject: Hole | $F_{(1,11)}=14.65$ | 0.0028* |
| | Between subject: Drug | $F_{(1,11)}=11.82$ | 0.0056* |
| 1F: OF distance | Unpaired t-test (Oxy, Sal) | $t_{(11)}=0.4254$ | 0.6787 |
| 1G: OF center (%) | Unpaired t-test (Oxy, Sal) | $t_{(11)}=0.3705$ | 0.7180 |
| 1H: SoP (%) | Unpaired t-test (Oxy, Sal) | $t_{(11)}=0.0554$ | 0.9568 |
| 1K: SuP (%) | Unpaired t-test (Oxy, Sal) | $t_{(11)}=0.7793$ | 0.4522 |
| 1L: EPM (%) | Unpaired t-test (Oxy, Sal) | $t_{(11)}=1.191$ | 0.2586 |
| **Figure 2: Early abstinence and late abstinence negative affect** |  |  |  |
| 2B: SA infusions | Two Way Mixed ANOVA |  |  |
| | Drug x Day interaction | $F_{(9,126)}=8.426$ | <0.0001* |
| | Within subject: Day | $F_{(3.210,44.94)}=8.384$ | 0.0001* |
| | Between subject: Drug | $F_{(1,14)}=7.969$ | 0.0136* |
| 2C: SA avg infusions | Mann-Whitney Test (Oxy, Sal) | U=0 | 0.0002* |
| 2D: SA avg pokes | Two Way Mixed ANOVA |  |  |
| | Hole x Drug interaction | $F_{(1,14)}=1.434$ | 0.2510 |
| | Within subject: Hole | $F_{(1,14)}=5.689$ | 0.0318* |
| | Between subject: Drug | $F_{(1,14)}=7.649$ | 0.0152* |
| 2E: Act pokes seeking | Two Way Mixed ANOVA |  |  |
| | AD x Drug interaction | $F_{(1,14)}=5.130$ | 0.0399* |
| | Within subject: AD | $F_{(1,14)}=33.12$ | <0.0001* |
| | Between subject: Drug | $F_{(1,14)}=10.76$ | 0.0055# |
| 2E: Inact pokes seeking | Two Way Mixed ANOVA |  |  |
| | AD x Drug interaction | $F_{(1,14)}=0.0092$ | 0.9248 |
| | Within subject: AD | $F_{(1,14)}=17.50$ | 0.0009* |
| | Between subject: Drug | $F_{(1,14)}=8.060$ | 0.0131* |
| 2F: Incubation score | Unpaired t-test (Oxy, Sal) | $t_{(14)}=2.265$ | 0.0399* |
| 2G: OF center (%) | Two Way Mixed ANOVA |  |  |
| | AD x Drug interaction | $F_{(1,14)}=1.711$ | 0.2119 |
| | Within subject: AD | $F_{(1,14)}=2.231$ | 0.1575 |
| | Between subject: Drug | $F_{(1,14)}=0.0008$ | 0.9781 |
| 2H: OF center entries | Two Way Mixed ANOVA |  |  |
| | AD x Drug interaction | $F_{(1,14)}=0.2423$ | 0.6302 |
| | Within subject: AD | $F_{(1,14)}=10.92$ | 0.0052* |
| | Between subject: Drug | $F_{(1,14)}=0.0937$ | 0.7641 |
| 2I: SoP (%) | Two Way Mixed ANOVA |  |  |
| | AD x Drug interaction | $F_{(1,14)}=0.1783$ | 0.6793 |
| | Within subject: AD | $F_{(1,14)}=11.98$ | 0.0038* |
| | Between subject: Drug | $F_{(1,14)}=0.9635$ | 0.3430 |
| 2J: SuP (%) | Two Way Mixed ANOVA |  |  |
| | AD x Drug interaction | $F_{(1,11)}=4.586$ | 0.0555# |
| | Within subject: AD | $F_{(1,11)}=5.135$ | 0.0446* |
| | Between subject: Drug | $F_{(1,11)}=3.850$ | 0.0756# |
| 2K: EPM (%) | Two Way Mixed ANOVA |  |  |
| | AD x Drug interaction | $F_{(1,14)}=0.0004$ | 0.9848 |
| | Within subject: AD | $F_{(1,14)}=8.208$ | 0.0125* |
| | Between subject: Drug | $F_{(1,14)}=0.5065$ | 0.4883 |
| **Figure 3: Housing condition and SA contribution to negative affect** |  |  |  |
| 3B: OF center (%) | Mann Whitney test (Single, Pair) | U=25 | 0.5054 |
| 3C: SoP (%) | Unpaired t-test (Single, Pair) | $t_{(14)}=1.378$ | 0.1899 |
| 3D: SuP (%) | Unpaired t-test (Single, Pair) | $t_{(14)}=2.420$ | 0.0297* |
| 3E: EPM center (%) | Mann Whitney test (Single, Pair) | U=20 | 0.2345 |
| 3F: OF center (%) | Kruskal-Wallis test (Single, Sal, Oxy) | H=0.4915 | 0.7821 |
| 3G: SoP (%) | One-Way ANOVA (Single, Sal, Oxy) | $F_{(2,34)}=1.906$ | 0.1642 |
| 3H: SuP (%) | One-Way ANOVA (Single, Sal, Oxy) | $F_{(2,31)}=5.204$ | 0.0113* |
| 3I: EPM (%) | Kruskal-Wallis test (Single, Sal, Oxy) | H=8.145 | 0.0170* |

Male and female rats underwent extended-access oxycodone or saline SA. The number of infusions across the 10 days of training is shown in Fig. 1B (drug x day interaction, F(_9,99_)=14.20, p<0.0001); nose pokes are shown in Fig. S1A. Oxycodone rats took more infusions than saline rats (Fig. 1C; unpaired t-test, t(_11_)=4.348, p=0.0012) and poked more in the active than inactive hole (Fig. 1D; hole x drug interaction, F(_1,11_)=5.589, p=0.0375) during the last 3 d of SA, indicating successful acquisition of oxycodone SA.

Weight loss typically occurs during non-contingent escalating dose opioid regimens and persists during early abstinence, (e.g. (11)). We recorded weight before SA training and on AD1-3. When data were normalized to weight before SA, oxycodone rats lost weight while saline rats gained weight over AD1-3 (Fig. 1E; AD x drug interaction, F(_3,33_)=23.90, p<0.0001), but there were no significant differences in absolute weight between the groups (Fig. S1B). After SA training, we assessed grooming, wet dog shakes, teeth chatters, jumps, and digging individually (Fig. S1), and we also calculated a cumulative score (Fig. 1F; see Methods). More teeth chatters (main effect of drug, F(_1,11_)=6.803, p=0.0243) and a trend towards more wet dog shakes (drug x AD interaction, F(_2,22_)=3.099, p=0.0652) were observed in oxycodone rats (Fig. S1C-G). No significant difference between oxycodone and saline rats was found when SWD symptoms were expressed as a cumulative score (Fig. 1F; AD x drug interaction, F(_2,22_)=1.597, p>0.05). There was no significant correlation between total oxycodone consumed (mg/kg) and the weight change or SWD symptoms (Table S2).

Starting on AD40, rats received a cue-induced seeking test followed by tests of negative affect (OFT, SoPT, SuPT, EPM), with the order of tests selected to minimize impact of earlier tests on later tests (Fig. 1A). Oxycodone rats showed greater cue-induced seeking than saline rats (Fig. 1G; main effect of drug, F(_1,11_)=11.82, p<0.0056) but subsequent tests for anxiety- or depressive-like behavior failed to reveal group differences. Specifically, there were no differences in distance traveled or time in center of an open field (Fig. 1H,I), preference for a stranger conspecific (Fig. 1J), sucrose preference (Fig. 1K), or time in open arms of the EPM (Fig. 1L) (unpaired t-tests, p>0.05). Additional measures made during these tests are shown in Fig. S1H-L.

Overall, these results suggest that our oxycodone SA regimen does not result in pronounced SWD signs in early abstinence (AD1-3) or negative affective states after protracted abstinence (AD41-46) compared to rats that self-administered saline.

### 3.2 Comparison of behaviors indicative of negative affect in early and late abstinence from saline or oxycodone self-administration

Although studies in Fig. 1 revealed no difference between saline and oxycodone rats in negative affect tests after protracted abstinence, we generated a second cohort of saline and oxycodone rats to determine if changes could be detected within each group as a function of abstinence duration. These rats received seeking tests and negative affect tests in both early abstinence (AD1-7) and protracted abstinence (AD40-46) (Fig. 2A). Full statistics for all results in Fig. 2 are shown in Table 1. As in the first cohort, oxycodone rats took more infusions than saline rats (Fig 2B; drug x day interaction, F(_9,126_)=8.426, p<0.0001; Fig 2C; Mann Whitney test, U=0, p=0.0002) and discriminated between active and inactive holes (Fig. 2D; main effect of hole, F(_1,14_)=5.689, p=0.0318). During seeking tests, responses in the previously active hole were significantly greater on AD40 than AD1 for oxycodone rats (Fig. 2E; Active pokes: AD x drug interaction, F(_1,14_)=5.130, p=0.0399) while inactive hole responses and active pokes by saline rats did not show significant time-dependency. Accordingly, the incubation score (AD40-AD1 active pokes) was significantly greater for oxycodone rats (Fig 2F; unpaired t-test, t(_14_)=2.265, p=0.0399). These data indicate that oxycodone rats demonstrated incubation of craving on AD40.

**Figure 2.**
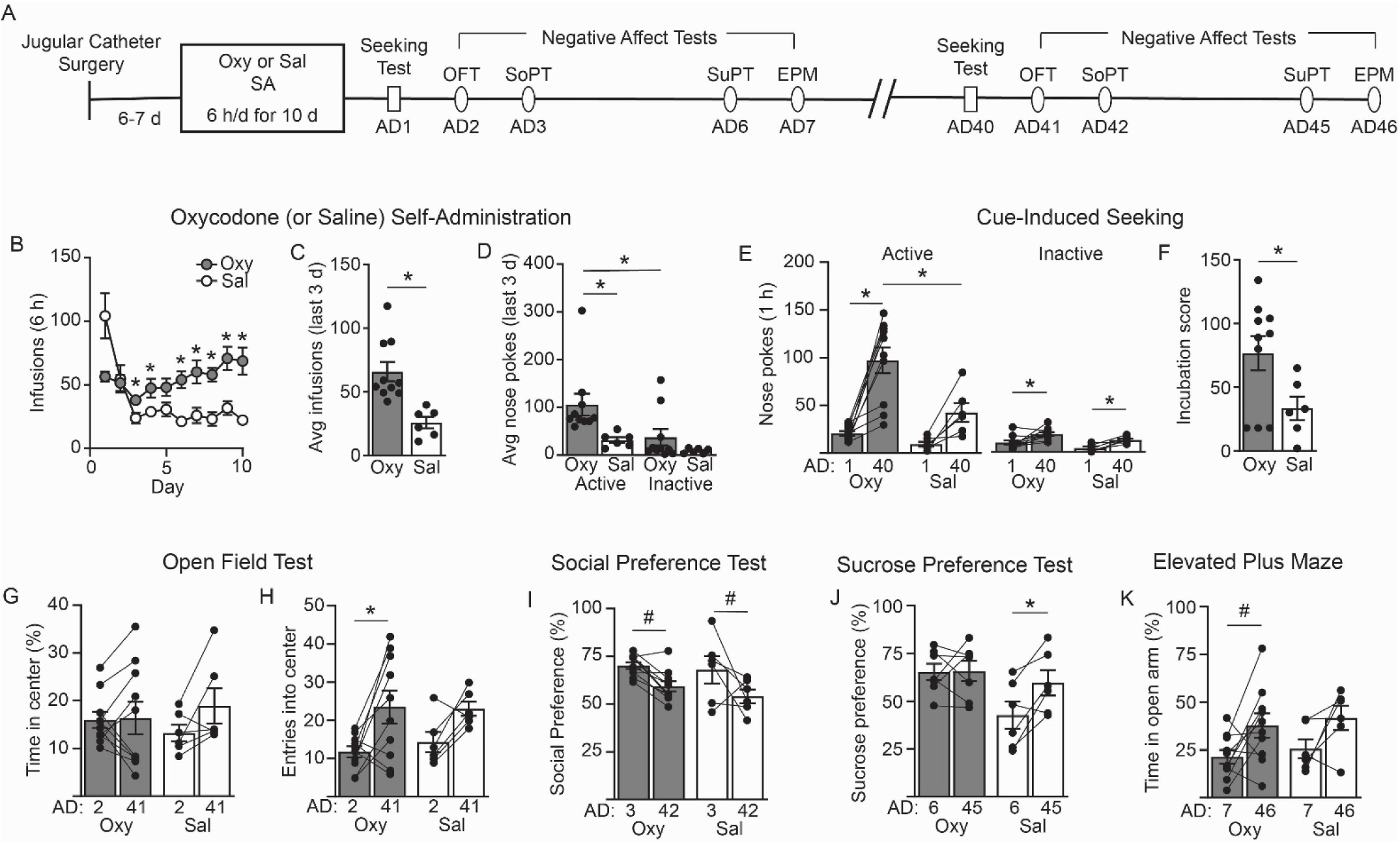
Negative affect during incubation of craving following oxycodone self-administration. A) Rats received jugular catheters and recovered for 6-7 days. Rats self-administered oxycodone (Oxy; 0.1 mg/kg/infusion; n=10) or saline (Sal; n=6) for 6 h/d for 10 d. Rats then underwent a series of behavior tests in early abstinence: cue-induced seeking tests (1 h) on abstinence day (AD) 1, open field test (OFT) on AD2, social preference test (SoPT) on AD3, sucrose habituation on AD4-5, sucrose preference test (SuPT) on AD6, and elevated plus maze (EPM) on AD7. Rats remained in their home cage until late abstinence and then underwent the same sequence of behavior tests on AD40-46. B) The number of infusions of Oxy or Sal that rats self-administered in 6 h across the 10 d of SA. Oxy rats took a larger number of infusions than Sal rats on days 3-4 and 6-10 (post hoc Sidak’s multiple comparisons test, p<0.05 for each day). C) During the last 3 d of SA, Oxy rats took a significantly higher number of infusions than Sal rats (Mann-Whitney Test, U=0, p=0.0002). D) During the last 3 d of SA, Oxy rats poked more in the active (Act) hole than the inactive (Ina) hole (post hoc Sidak’s multiple comparisons test, p=0.0220), an effect not observed in Sal rats. Oxy rats poked more in the active hole than Sal rats (post hoc Sidak’s multiple comparisons test, p=0.0186). E) The number of active and inactive pokes during cue-induced seeking tests on AD1 and AD40. Oxy rats poked more in the active hole on AD40 than on AD1 (post hoc uncorrected Fisher’s LSD, p<0.0001) and compared to Sal rats (post hoc uncorrected Fisher’s LSD, p=0.0005). Both Oxy and Sal rats show an increase in poking in the inactive hole from AD1 to AD40 (post hoc Sidak’s multiple comparisons test, Oxy p=0.0071, Sal p=0.0428). F) The incubation score (AD40 Act-AD1 Act) was significantly higher in Oxy rats compared to Sal rats (unpaired t-test, p=0.0399). G) Oxy and Sal rats spent the same amount of time in the center of the open field on AD2 and AD41. H) Oxy rats entered the center of the open field more times on AD41 compared to AD1 (post hoc Sidak’s multiple comparisons test, p=0.0156). I) Oxy and Sal rats trended towards a reduction in time spent exploring a novel conspecific from AD3 to AD42 (post hoc Sidak’s multiple comparisons test AD3 v AD42, Oxy: p=0.0521, Sal: p=0.0547). J) Sal rats increased sucrose preference from AD6 to AD45 (post hoc Sidak’s multiple comparisons, p=0.0239) whereas Oxy rats did not change. K) Oxy rats trended towards an increase in the amount of time spent in the open arm from AD7 to AD46 (post hoc Sidak’s multiple comparisons test, p=0.0661). *p<0.05, #p<0.07. Abbreviations: abstinence day (AD), average (avg), elevated plus maze (EPM), open field test (OFT), oxycodone (Oxy), saline (Sal), self-administration (SA), social preference test (SoPT), sucrose preferencetest (SuPT). Full statistical results for Fig. 2 are provided in Table 1. See Fig. S2 and Table S1 for additional analyses.

Interestingly, both saline and oxycodone rats showed some time-dependent changes in negative affect tests. Commonly reported metrics are shown in Fig. 2, while Fig. S2 shows additional supporting analyses (full statistics for Fig. S2 in Table S1). In OFT tests, total distance traveled (Fig. S2B) and time in the center (Fig. 2G) did not differ as a function of abstinence day or drug group. However, entries into the center of the open field (Fig. 2H) increased from AD2 to AD41 (main effect of AD, F(_1,14_)=10.92, p=0.0052), with post hoc tests indicating a significant effect for oxycodone rats (p=0.0156). In the SoPT, time spent with the novel conspecific rat (Fig. 2I) showed a significant main effect of AD (Table 1; F(_1,14_)=11.98, p=0.0038) with post hoc tests showing trends towards lower preference on AD42 for both saline and oxycodone rats compared to AD3 (p=0.0547 and 0.0521, respectively). Interestingly, entries into the social zone (Fig. S2D) tended to increase in late abstinence. In the SuPT, saline rats showed greater preference on AD45 vs AD6 while oxycodone rats did not change (Fig. 2J; main effect of AD, F(_1,11_)=5.135, p=0.0446). In EPM tests, while entries into the open arm did not differ by time or group (Fig. S2F), time spent in the open arm (Fig. 2K) showed a main effect of AD (F(_1,14_)=8.208, p=0.0125) with post hoc tests showing that oxycodone rats trended towards more time in the open arm (p=0.0661).

These results suggest that modest time-dependent changes occur in both groups, although they are not suggestive of intensification of negative affect in oxycodone rats as abstinence progresses. Thus, oxycodone rats show less anxiety-like behavior in late abstinence in some tests (center entries in OFT, open arm time in EPM) although saline rats showed similar trends (Figs. 2H and 2K). Both oxycodone and saline rats trend towards lower social preference in late abstinence (Fig. 2I) and saline rats show a small increase in sucrose preference in late abstinence (Fig. 2J). However, when saline and oxycodone rats are compared in late abstinence, data in both Figs. 1 and 2 indicate no robust differences between the groups.

### 3.3 Comparison of single and pair-housed rats

During incubation experiments, rats are single-housed for ∼9 weeks (Figs. 1A & 2A). To determine if this influences our measures of negative affect, we compared drug-naïve rats that were single-or pair-housed for a similar length of time (56-61 days; Fig. 3A) using OFT, SoPT, SuPT, and EPM (Figs. 3B-E, Table 1). The only group difference identified was lower sucrose preference in pair-housed rats (Fig. 3D; unpaired t-test, t(_14_)=2.42, p=0.0297). Additional metrics are shown in Fig. S3 (total distance traveled and entries into center for OFT; time in social zone and entries into social zone for SoPT; amount consumed for SuPT; entries into open arm for EPM; statistical results in Table S1). Figs. 3F-I show data from the drug-naïve single-housed rats replotted alongside oxycodone and saline forced abstinence groups (combined from Figs. 1 and 2). No group differences were found for time in center of the open field or social preference (Figs. 3F,G) but single-housed, drug-naïve rats showed greater sucrose preference (Fig. 3H; One-Way ANOVA, F(_2,31_)=5.204, p=0.0113) and time in the open arm of the EPM (Fig. 3I; Kruskal-Wallis test, H=8.145, p=0.0170) compared to saline and oxycodone groups (which did not differ from each other).

**Figure 3.**
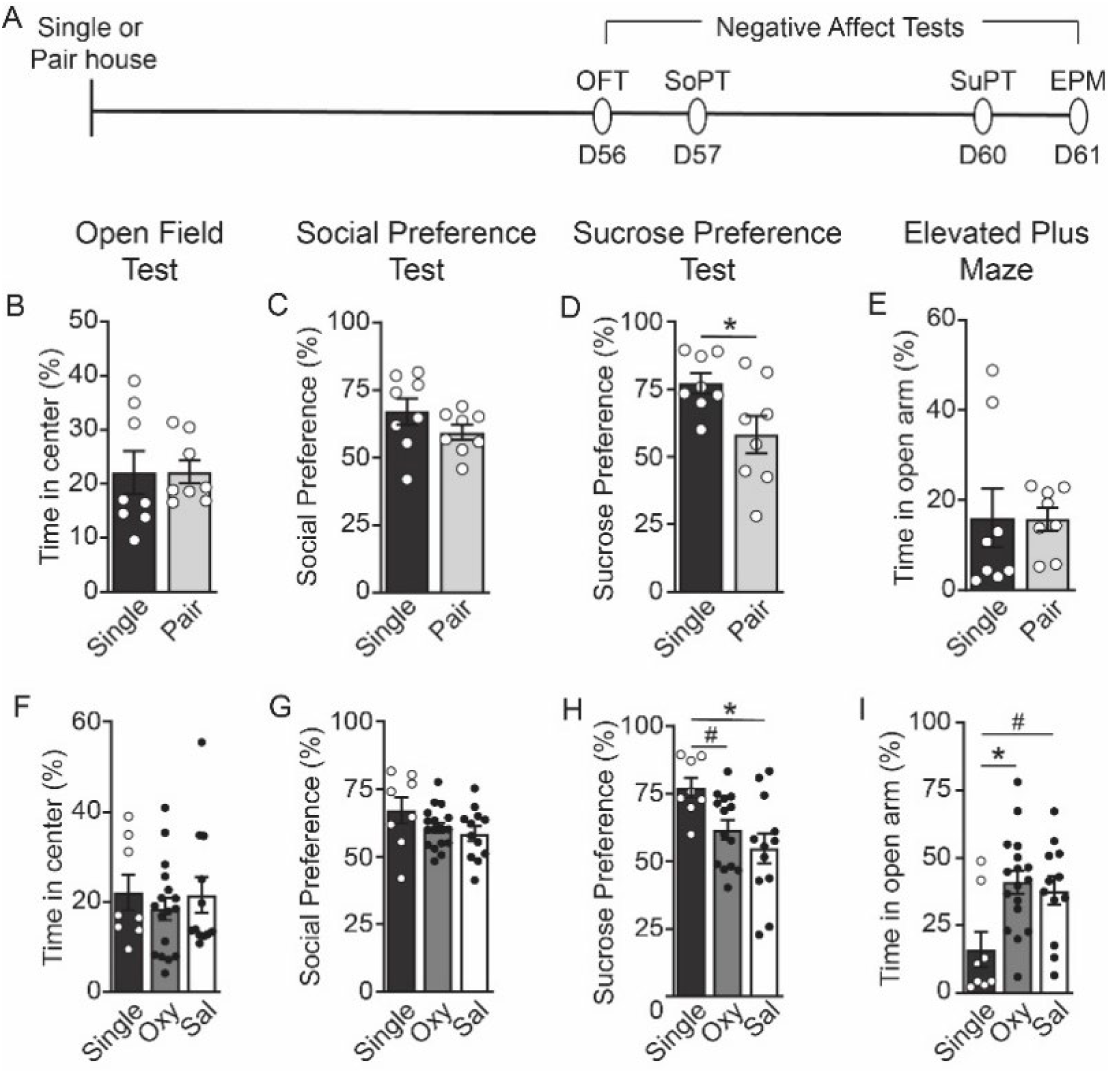
Effects of housing condition and self-administration training on assays measuring negative affect. A) A cohort of naïve rats were single- or pair-housed for 8 weeks. Rats then underwent open field test (OFT) on day 56, social preference assay test (SoPT) on day 57, sucrose preference test (SuPT) on days 58-60, and elevated plus maze (EPM) on day 61. B) Single-housed and pair-housed rats spent the same amount of time in the center of the open field. C) The preference for a novel rat was no different between single- or pair-housed rats. D) Pair-housed rats exhibited a lower sucrose preference than single-housed rats (unpaired t-test, p=0.0399). E) Housing status did not change the amount of time spent in the open arm of the EPM. F) Rats that self-administered oxycodone (Oxy) or saline (Sal) spent a similar amount of time in the center of the open field as single-housed, drug-naïve rats. G) Self-administration did not alter the time spent exploring a novel rat compared to single-housed rats. H) While Sal and Oxy rats did not demonstrate differences in sucrose preference, single-housed rats had higher sucrose preference compared to Sal (post hoc Tukey’s multiple comparisons test p=0.0086) and Oxy (p=0.0736) rats. I) While Sal and Oxy rats did not demonstrate differences in the time spent in the open arm of the EPM, single-housed rats spent less time in the open arm of the EPM compared to Sal (post hoc Dunn’s multiple comparisons test, p=0.0559) or Oxy (p=0.0181) rats. *p<0.05, #p<0.08. Abbreviations: elevated plus maze (EPM), open field test (OFT), oxycodone (Oxy), saline (Sal), social preference test (SoPT), sucrose preference test (SuPT). Full statistical results are provided in Table 1. See Fig. S3 and Table S1 for additional analyses.

Together, these results suggest increased anhedonia-like behavior (SuPT) and reduced anxiety-like behavior (EPM) in saline and oxycodone rats after protracted abstinence compared to drug-naïve rats singly housed for a similar duration, possibly related to a history of surgery and operant training in the former groups.

## 4. Discussion

### 4.1 Overview

Our results support the following conclusions, each of which will be discussed below. First, although we used an oxycodone SA regimen that elicits robust incubation of cue-induced oxycodone seeking (and is very similar to regimens used by other labs; see Introduction), this incubation was not accompanied by anxiety- or depressive-like behavior as compared to control rats that self-administered saline. Our oxycodone regimen also failed to elicit pronounced spontaneous SWD signs during AD1-3. Second, while we identified modest changes in affective states in both saline and oxycodone rats when comparing early and late abstinence timepoints within each treatment group, these changes did not support the idea that progressive increases in negative affective state accompany the emergence of incubation of craving. Third, studies of drug-naïve rats that were single-housed or pair-housed for a duration of time similar to single-housing during an incubation experiment (8+ weeks) did not show differences in affective state.

Regarding sex as a biological variable, we used males and females in all experiments and did not observe obvious differences. Because we (22) and others (3, 31) have previously found similar incubation of opioid craving in male and female rats (but see (25)) and because sex was not the primary independent variable of interest in our behavioral studies, they were powered to detect effects of treatment but not sex. Nonetheless, we acknowledge that this is a limitation of our study in light of sex differences in other behaviors related to oxycodone SA in rodents (26, 32–35) and in human studies (36, 37). However, the only paperexamining behaviors related to negative affect after oxycodone SA found no sex differences in these behaviors (32).

### 4.2 Negative affective states do not emerge in rats exhibiting incubation of oxycodone craving

It is well established that people recovering from OUD often experience persistent negative affective states that contribute to relapse and that negative affective states are observed in rodents after non-contingent escalating dose regimens of opioids and can sometimes intensify over several weeks of withdrawal (see Introduction). Far less is known about whether negative affective states occur after protracted abstinence from opioid self-administration. In our first cohort of rats, we performed OFT, SoPT, SuPT, and EPM after protracted withdrawal (AD40+) from oxycodone SA and observed no differences from saline controls. No other studies have assessed negative affect after protracted oxycodone withdrawal, but two have done so after heroin SA. First, a study of incubation of heroin craving measured ultrasonic vocalizations (USV) during AD1 and AD30 seeking tests; they found an increase in 50-kHz USVs (which indicate a rewarding situation) on WD30 in males but not females but detected very few 22-kHz USVs (which indicate a negative state) in any group (38). Relatively low 22-kHz USVs after heroin incubation are consistent with our findings of no obvious negative affective states after oxycodone incubation. Second, a recent study found that males and female mice tested 14 days after discontinuing heroin self-administration spent significantly less time interacting with a social stimulus (age- and sex-matched mouse) compared to saline controls (39). In mice with lower than median heroin seeking on AD13 there was a significant negative correlation between social interaction time and heroin seeking (39). In this prior study, only the social stimulus was present during the test. We used a different social interaction test in which rats could interact with a stranger conspecific or an empty container during the test and results are reported as percentage of time interacting with the stranger conspecific (see Methods) and found no difference between saline and oxycodone rats (Figs. 1 and 2). The use of different test procedures, as well as different opioids and species, may underlie the different results.

While we did not assess negative affect in early abstinence in this cohort of rats, we quantified spontaneous SWD signs on AD1-3. While oxycodone rats showed modest increases in teeth chatters and wet dog shakes, we observed no significant differences in the cumulative somatic withdrawal score between saline and oxycodone rats. In a prior study using a short-access oxycodone SA regimen, SWD signs were reported 0 and 22 h after the last session but their magnitude is difficult to assess because vehicle controls were not included (27). Another study found significant SWD signs in oxycodone rats compared to vehicle rats 24 h after discontinuing an extended-access regimen that included more sessions than ours (22 vs 10) (40). Notably oxycodone rats in our study exhibited reduced body weight on AD1-3, consistent with prior reports using non-contingent morphine dosing (11), suggesting that physiological signs of withdrawal may be present even in the absence of robust expression of SWD signs. Furthermore, we did not assess naloxone-precipitated withdrawal. Thus, we cannot exclude the possibility that physical dependence was present but not detectable under our conditions.

Two factors likely contribute to lack of pronounced SWD signs and negative affect in our self-administration study relative to non-contingent escalating dose regimens (see Introduction). First, volitional drug intake may be less aversive or engage distinct motivational or neurobiological processes compared to experimenter-administered drugs. Second, total opioid exposure is lower in self-administration studies. In the present study, rats (300-400 g) self-administered 15-20 mg of oxycodone over 10 days, whereas non-contingent escalating regimens deliver substantially higher doses over shorter timeframes (e.g., ∼63 mg oxycodone in a 20-25 g mouse over 9 days (41)), resulting in markedly greater exposure relative to body mass.

### 4.3. Time-dependent changes in negative affective behavior over 40+ days of abstinence

Although the experiment shown in Fig. 1 did not detect negative affective states after protracted withdrawal from oxycodone SA, this experiment involved a between-group (saline/oxycodone) comparison at a single range of abstinence times (AD41-46). We therefore generated a second cohort of saline and oxycodone SA rats for a within-subject study comparing early (AD2-7) and late (AD41-46) abstinence. As detailed in Results and in Fig. 2, we did find some time-dependent changes within each treatment group. Overall, however, the measured behaviors in saline rats remained stable across abstinence with only a modest reduction in social preference and increase in sucrose preference in late abstinence, whereas oxycodone rats showed a similar reduction in social preference but also exhibited OFT and EPM behaviors consistent with reduced anxiety-like behavior in late versus early abstinence. It is worth noting that while many studies report reduced time in the open arm of the EPM after escalating morphine doses, including two conducted after 1-4 weeks of withdrawal (16, 42), one study found the opposite result (similar to ours) in male mice 6 weeks after morphine withdrawal and interpreted this as impulsive behavior (43), which has been observed in human opioid users (e.g., (44)). Overall, our results do not support the idea that negative affective states intensify as abstinence progresses and contribute to the incubation of oxycodone craving through a negative reinforcement mechanism. Likewise, a study of heroin incubation discussed in the previous section found no change in 22-kHz USVs (which indicate a negative state) over the course of heroin incubation (38).

The absence of differences in negative affect between saline and oxycodone rats in early abstinence is interesting to consider in light of prior studies. Focusing first on oxycodone, tests performed in the first 24 h after an oxycodone SA session revealed irritability(32, 45) and elevated brain reward threshold (46), but these studies used a higher oxycodone dose (0.15 mg/kg/infusion) and a longer daily session (12-h). A 12-h session was also used in a study reporting potentiated acoustic startle, interpreted as anxiety-like behavior, after 10-20 h of withdrawal from heroin SA (47).

### 4.4. Single-housing alone does not necessarily elicit negative affective states

We evaluated the impact of single-housing in drug-naïve rats to determine if post-surgical isolation during incubation-style abstinence experiments might influence measures of negative affect. Under our conditions, single-versus pair-housing for a duration comparable to protracted abstinence did not alter performance in OFT, SoPT, or EPM although pair-housed rats showed increased anhedonia-like behavior (SuPT). Notably, when compared to these single-housed drug-naive rats, rats that underwent either oxycodone or saline self-administration exhibited increased anhedonia-like behavior (SuPT) and reduced anxiety-like behavior (EPM). Overall, these results suggest that prior surgical and operant training experience had a greater impact on affective state than housing conditions. Prior work has shown that the effects of social isolation on affective states are variable, with minimal impact on locomotion and unconditioned anxiety-like behaviors (i.e., OFT and EPM) and moderate effects on sucrose preference and social preference tests (48, 49). Furthermore, the effects of social isolation depend on factors such as sex, strain, and comparator housing conditions. In contrast to prior studies that typically initiate isolation during adolescence (e.g., (48, 49)), our study examined isolation beginning in young adulthood for a duration aligned with incubation procedures. These differences in developmental timing and protocol are likely to influence the emergence of affect-related behaviors. Collectively the available evidence suggests that single-housing rats is not sufficient to induce negative affect-like behavior, particularly in standard colony environments where rats maintain auditory, visual and olfactory interactions with conspecifics in the housing room. We also note that single vs. pair housing did not affect incubation of heroin craving in either sex (50).

## 5. Conclusions

We conclude that negative affective states do not occur alongside incubation of craving using a standard oxycodone incubation regimen, probably due in part to lower drug intake during our SA regimen compared to non-contingent escalating dose opioid regimens typically used to study SWD signs and negative affective behavior. This does not diminish the utility of the incubation model for studying cue-induced opioid craving and its neurobiological basis. Furthermore, we note that the level of oxycodone intake in our paradigm approximates clinically relevant exposure associated with pain management in humans (see Table S3), raising the possibility that clinically relevant patterns of opioid intake may elicit persistent adaptations that increase reactivity to opioid-related cues.

## Author Contributions

Conception & Study Design: AMW, KAM, MEW. Data Acquisition: AMW, AG, ABK, JW, HMK, MMB. Analysis and/or Interpretation of Data: AMW, MEW. Drafting of manuscript: AMW, MEW. Obtained funding: MEW. All authors reviewed and approved the manuscript.

## Funding Statement

This work was supported by USPHS grant DA059601 (MEW). KAM was supported by a predoctoral NSF fellowship and HMK by F31 DA064365. KAM and HMK also received ARCS Foundation Oregon Scholarships.

## Ethics Statement

Animal studies were approved by the Institutional Animal Care and Use Committee at Oregon Health & Science University in accordance with the NIH Guide for the Care and Use of Laboratory Animals.

## Data availability

Data are available upon request from the communicating author.

## Conflict of Interest Statement

The authors declare no conflict of interests.

## Supplemental Material (Figures S1-S3, Tables S1-S3)

**Figure S1.**
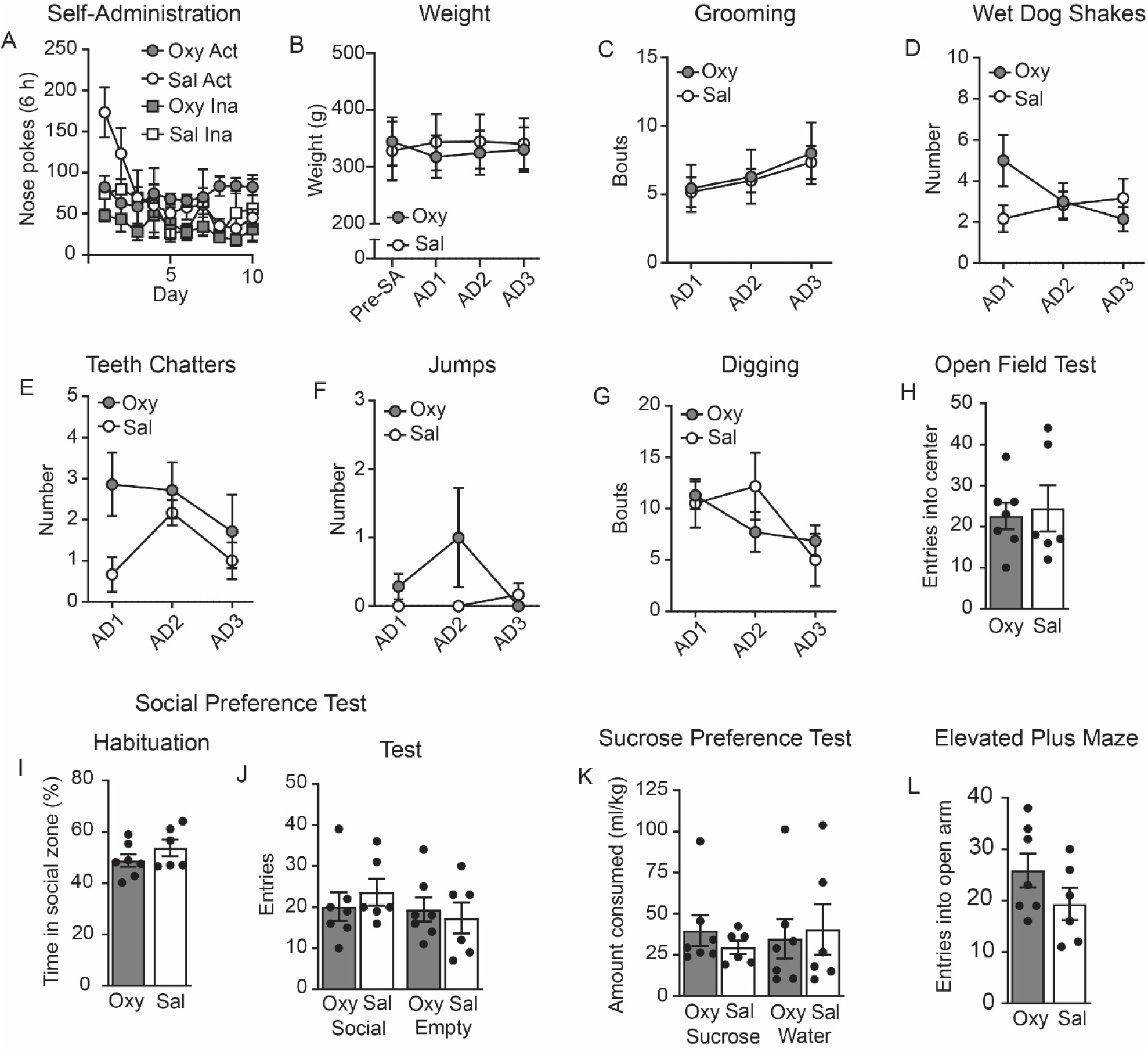
Additional drug self-administration, somatic withdrawal, open field, social preference test, sucrose preference test, and elevated plus maze data associated with. Figure 1. Full statistical analysis for data presented in this figure is shown in Table S1. A) Nose pokes in the active (Act) and inactive (Ina) hole in rats that self-administered (SA) oxycodone (Oxy) or saline (Sal). B) The weight of rats that self-administered Oxy did not differ from rats that self-administered Sal before SA or on AD1-3. C) The number of grooming bouts was the same in Oxy and Sal rats on AD1-3. D) Oxy and Sal rats exhibited the same number of wet dog shakes on AD1-3. E) The number of teeth chatters expressed by Oxy and Sal rats on AD1-3 were the same. F) Oxy and Sal rats jumped the same number of times on AD1-3. G) Oxy SA did not alter the number of digging bouts on AD1-3 compared to Sal rats. H) Oxy and Sal rats entered the center of the open field the same number of times. I) Prior to social preference test, Oxy and Sal rats spent the same amount of time in the zone where the novel conspecific was placed. J) During the social preference test session, Oxy and Sal rats made the same number of entries into the chamber containing a novel conspecific as the chamber without a rat. K) The amount of sucrose and water consumed (ml/kg) was the same between Oxy and Sal rats. L) The number of entries into the open arm was the same between Oxy and Sal rats. Abbreviations: abstinence day (AD), active (Act), average (avg), inactive (Ina), oxycodone (Oxy), saline (Sal), self-administration (SA).

**Figure S2.**
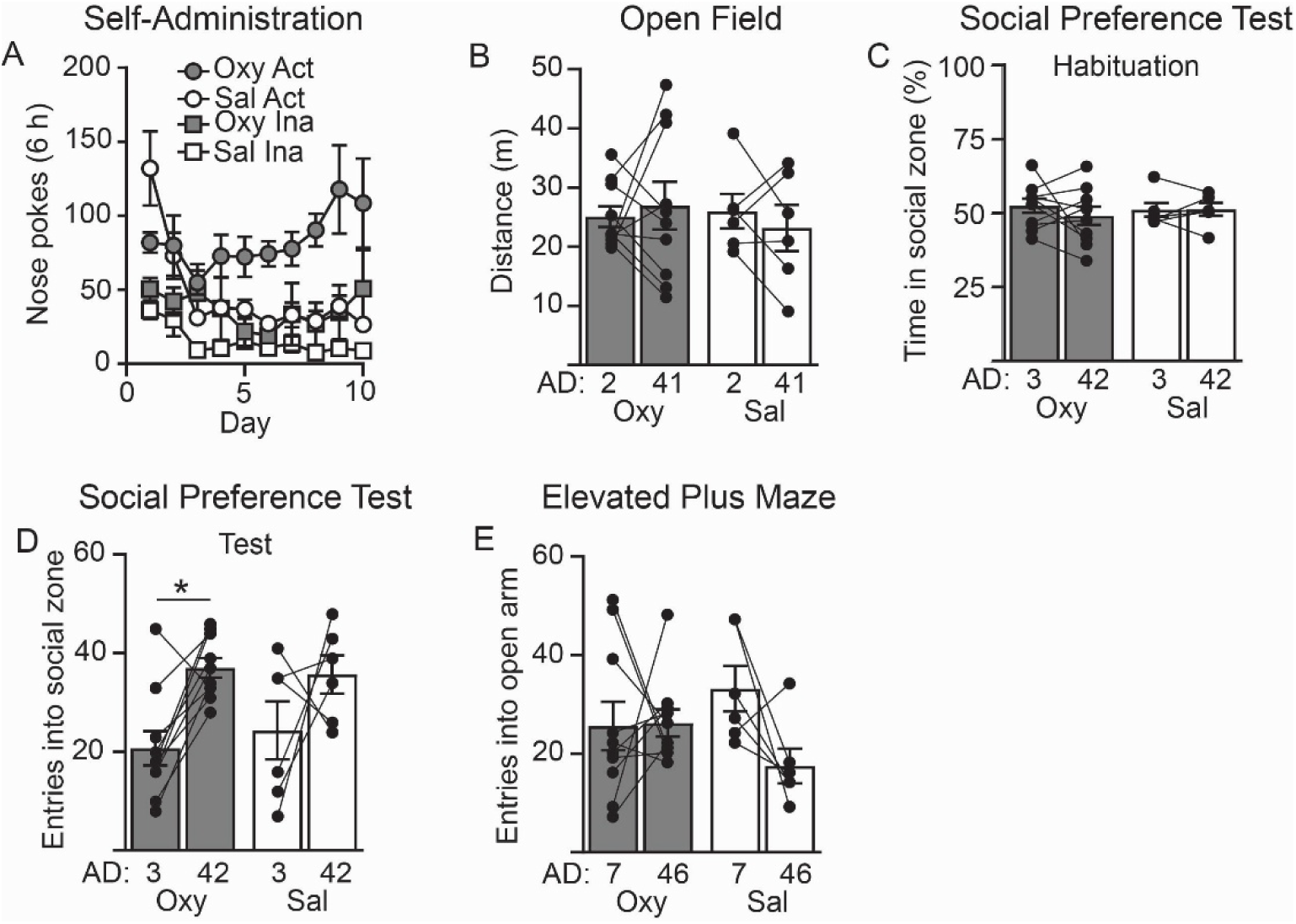
Additional self-administration, open field, social preference test, sucrose preference test, and elevated plus maze data associated with. Figure 2. Full statistical analysis for data presented in this figure is shown in Table S1. A) Nose pokes in the active (Act) and inactive (Ina) hole in rats that self-administered oxycodone (Oxy) or saline (Sal). B) Both Oxy and Sal rats travelled the same distance in the open field on abstinence day (AD) 2 and AD41. C) Prior to testing, Oxy and Sal rats did not exhibit a preference for the zone where the novel conspecific was placed. D) Oxy rats entered the zone around the novel conspecific more on AD42 compared to AD3 (post hoc Sidak’s multiple comparisons test, p=0.0149) whereas this was not significant for Sal rats. E) Neither group showed a significant difference in entries into the open arm on AD7 versus AD46. *p<0.05. Abbreviations: abstinence day (AD), active (Act), inactive (Ina), oxycodone (Oxy), saline (Sal).

**Figure S3.**
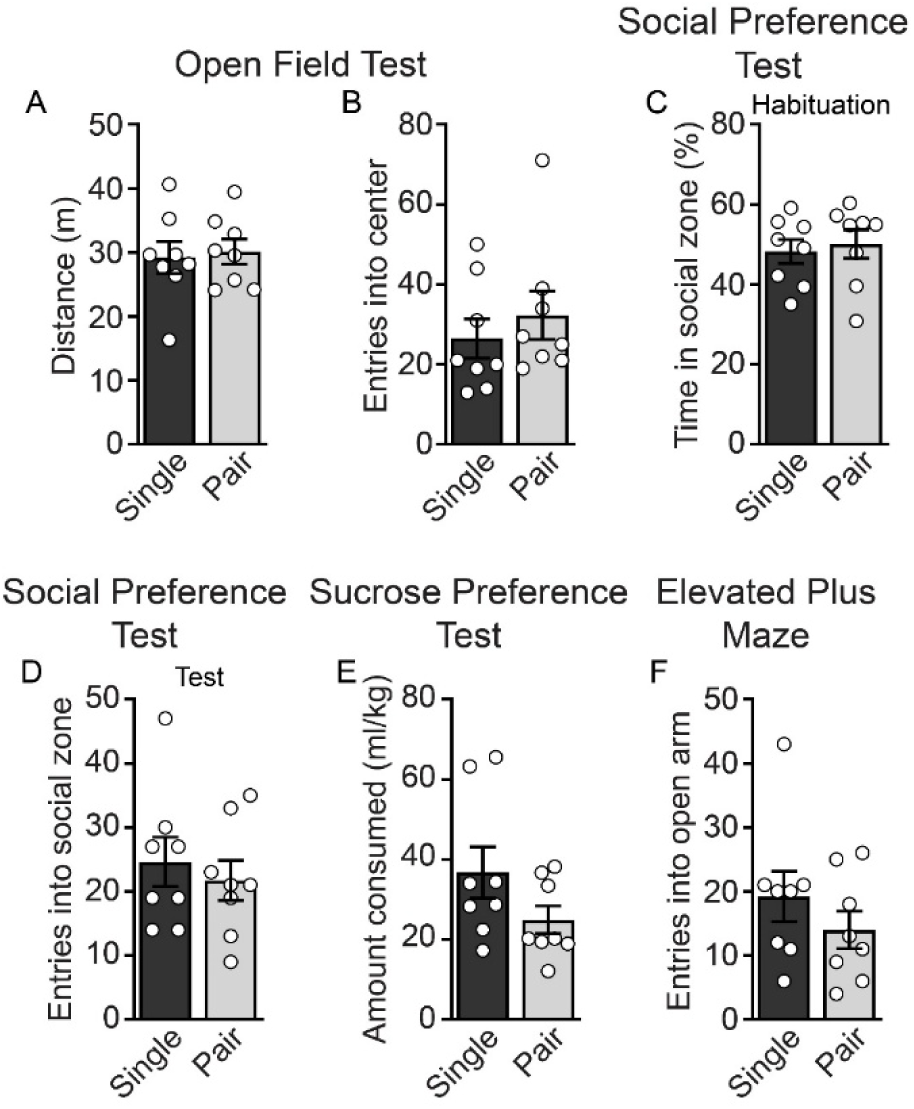
Additional open field test, social preference test, sucrose preference test, and elevated plus maze data associated with. Figure 3. Full statistical analysis for data presented in this figure is shown in Table S1. A-B) The total distance traveled (A) and number of entries in the center (B) of the open field were not different between single-housed or pair-housed rats. B-C) Single-housed and pair-housed rats did not exhibit a preference for either side of the social assay chamber during habituation (C) and entered the zone containing the novel conspecific rat the same number of times (D) during the social preference test. E) The amount of sucrose consumed during the sucrose preference test was the same between single-housed and pair-housed rats. F) Single-housed and pair-housed rats entered the open arm on the elevated plus maze the same number of times.

**Table S1.**
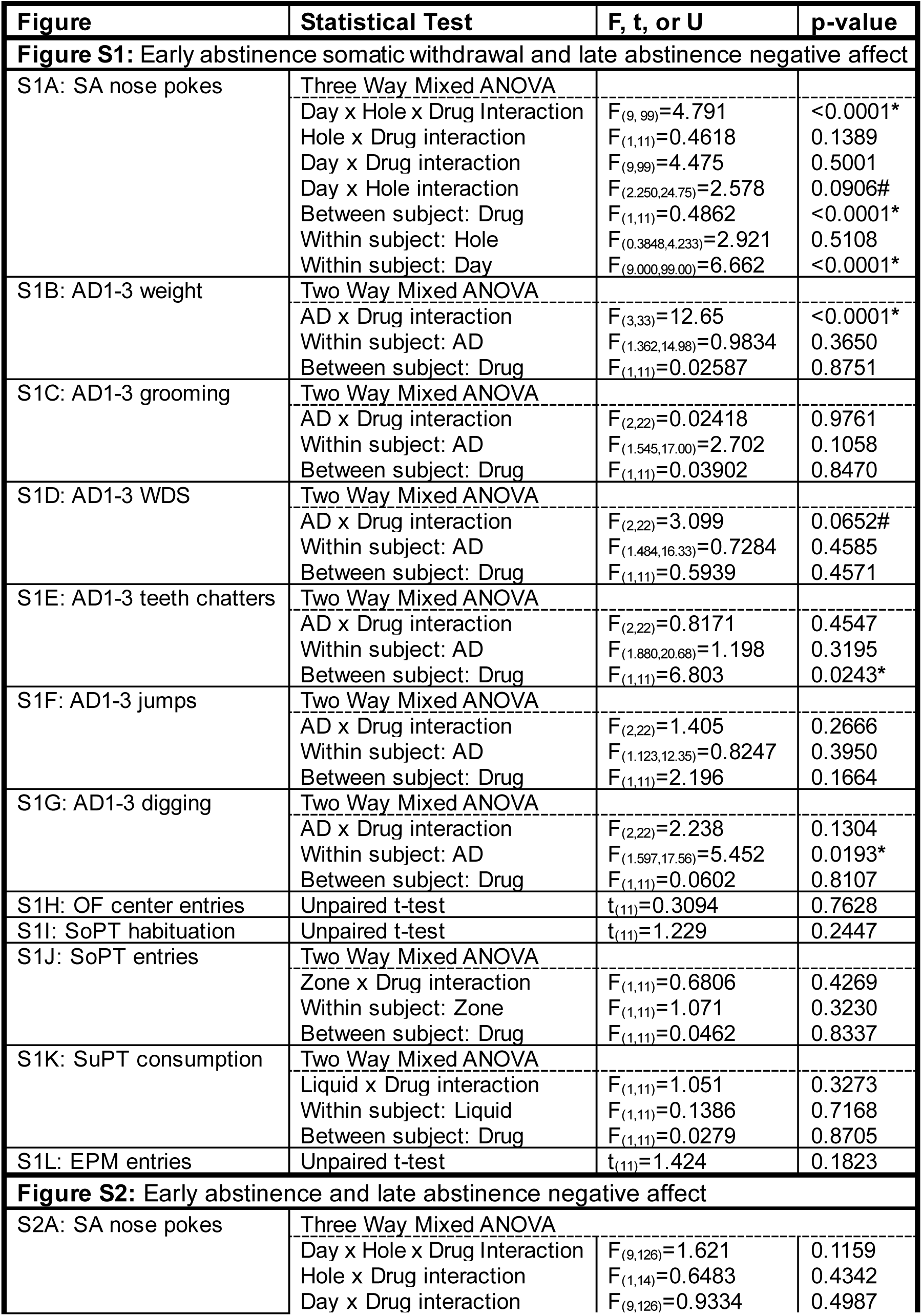
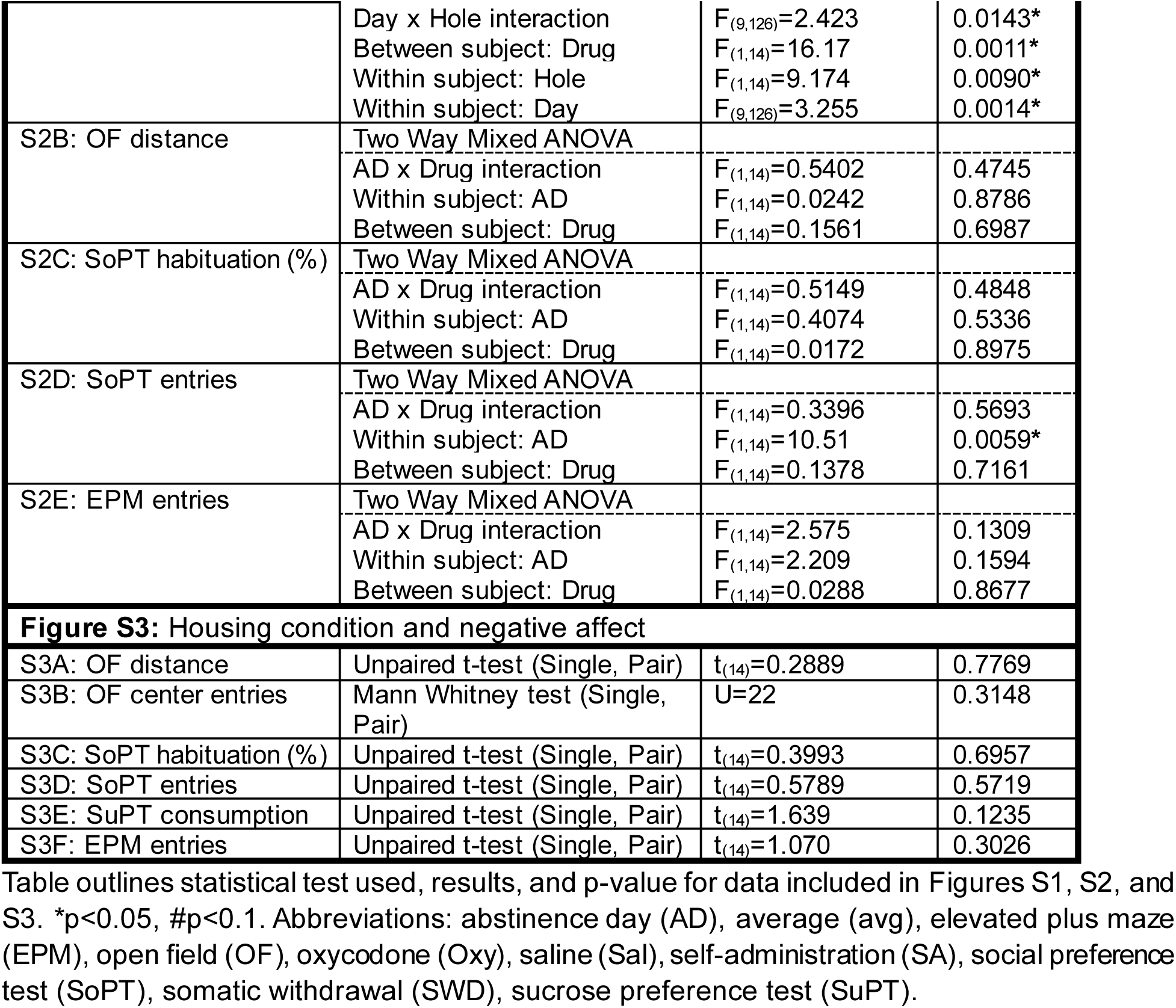
Full statistical results for all data presented in Supplementary Figures.

**Table S2.** Correlation analysis of oxycodone (Oxy) consumption and spontaneous withdrawal signs.

| **Variable 1** | **Variable 2** | **n** | **Pearson's r** | **p** |
| --- | --- | --- | --- | --- |
| Oxy consumed (mg/kg) | Weight change | 7 | 0.04249 | 0.928 |
| Oxy consumed (mg/kg) | Somatic withdrawal score | 7 | 0.5462 | 0.205 |
| Oxy consumed (mg/kg) | Grooming bouts | 7 | 0.1721 | 0.712 |
| Oxy consumed (mg/kg) | Wet dog shakes | 7 | 0.7054 | 0.077 |
| Oxy consumed (mg/kg) | Teeth chatters | 7 | 0.4680 | 0.290 |
| Oxy consumed (mg/kg) | Jumps | 7 | -0.1357 | 0.772 |
| Oxy consumed (mg/kg) | Digging bouts | 7 | 0.5931 | 0.160 |

**Table S3.** Rat equivalent daily dosing of oxycodone (Oxy) relative to human dosing for pain management

| Human Oxy Dosing | Daily Human Oxy Intake | K<sub>m</sub> (for rats) | Rat Oxy Equivalent Dose | Rat Oxy Dosing | Average Oxy Intake during SA |
| --- | --- | --- | --- | --- | --- |
| 20-60 mg per day | 0.33-1 mg/kg | 6.2 | 2.05-6.2 mg/kg | 0.41-2.79 mg per day | 1.71 mg per day |

Table S3 summarizes the conversion of clinically relevant human oxycodone (Oxy) dosing to rat-equivalent doses using allometric scaling (1). Human dosing was based on a typical pain management regimen of 5–15 mg administered every 6 hours (20–60 mg/day), corresponding to 0.33–1 mg/kg/day assuming a 60 kg individual. Rat-equivalent doses were calculated by multiplying the human mg/kg dose by the rat K<sub>m</sub> factor (6.2), yielding 2.05–6.2 mg/kg/day. This was converted to a total daily dose using an average rat body weight of 0.2-0.45 kg (0.41–2.79 mg/day). Average oxycodone intake during self-administration was calculated from infusion data in this study (mean=52.6 infusions/day) and average 0.325 kg body weight, resulting in an average intake of 1.71 mg/day. Notably, this level of intake falls within the range of rat-equivalent doses derived from clinically relevant human pain management regimens.

## Notes

### Competing Interest Statement

The authors have declared no competing interest.

